# Practice induces a qualitative change in the memory representation for visuomotor learning

**DOI:** 10.1101/226415

**Authors:** David M. Huberdeau, John W. Krakauer, Adrian M. Haith

**Affiliations:** Department of Biomedical Engineering, Johns Hopkins University School of Medicine, Baltimore, MD 21287.; Department of Neurology, Johns Hopkins University School of Medicine, Baltimore, MD 21287.; Department of Neuroscience, Johns Hopkins University School of Medicine, Baltimore, MD 21287.

**Author notes:** Equal contributions (John W. Krakauer, Adrian M. Haith).

## Abstract

Adaptation of our movements to changes in the environment is known to be supported by multiple learning processes which act in parallel. An implicit process recalibrates motor output to maintain alignment between intended and observed movement outcomes (“implicit recalibration”). In parallel, an explicit learning process drives more strategic adjustments of behavior, often by deliberately aiming movements away from an intended target (“deliberate re-aiming”). It has long been established that people form a memory for prior experience adapting to a perturbation through the fact that they become able to more rapidly adapt to familiar perturbations (a phenomenon known as “savings”). Repeated exposures to the same perturbation can further strengthen savings. It remains unclear, however, which underlying learning process is responsible for this practice-related improvement in savings. We measured the relative contributions of implicit recalibration and deliberate re-aiming to adaptation during multiple exposures to an alternating sequence of perturbations over two days. We found that the implicit recalibration followed an invariant learning curve despite prolonged practice, and thus exhibited no memory of prior experience. Instead, practice led to a qualitative change in re-aiming which, in addition to supporting savings, became able to be expressed rapidly and automatically. This qualitative change appeared to enable participants to form memories for two opposing perturbations, overcoming interference effects that typically prohibit savings when learning multiple, opposing perturbations. Our results are consistent with longstanding theories that frame skill learning as a transition from deliberate to automatic selection of actions.

## Introduction

Motor learning is often studied using adaptation tasks (Cunningham, 1989; Kluzik et al., 2008; Krakauer et al., 1999, 2000; Martin et al., 1996a; Shadmehr and Mussa-Ivaldi, 1994; Welch et al., 1993; Wolpert et al., 1995). In these tasks, a systematic perturbation is applied during a movement, and participants must learn to adjust their actions to cancel the effects of the perturbation and regain baseline levels of performance. The ability to adapt to an imposed perturbation appears to be supported by at least two underlying learning processes (Huberdeau et al., 2015a). One is an implicit recalibration process which is known to be cerebellum-dependent and driven by sensory prediction errors (Mazzoni and Krakauer, 2006; Taylor et al., 2010). The second process is deliberate compensation for imposed perturbations through re-aiming of their reaching movements (Fernandez-Ruiz et al., 2011; Haith et al., 2015; Martin et al., 1996a; Morehead et al., 2015; Taylor et al., 2014).

Experiencing a particular perturbation multiple times usually results in successful adaptation in fewer trials, a phenomenon known as *savings* (Brashers-Krug et al., 1996; Ebbinghaus, 1913; Huang et al., 2011, 2011; Kojima et al., 2004; Lackner and Lobovits, 1977; Villalta et al., 2013; Zarahn et al., 2008). Reach directions typically revert to baseline with time or when feedback is removed (Joiner and Smith, 2008; Kitago et al., 2013), leaving savings as the most reliable sign of long-term memory for prior adaptation. A number of recent studies have shown that savings is not attributable to faster implicit recalibration (as suggested by (Herzfeld et al., 2014)), but is instead solely attributable to more effective deliberate re-aiming (Hadjiosif and Smith, 2013; Haith et al., 2015; Morehead et al., 2015), likely through retrieval of a previously successful re-aiming strategy (Haith and Krakauer, 2014; Huberdeau et al., 2015b). These results suggest, puzzlingly, that long-term memory for adaptation is represented as a memory that must be expressed deliberately, rather than as a motor memory, which we would expect to be expressed automatically.

This conclusion that savings is deliberate seems incongruent with everyday experience acting under perturbed mappings (e.g. wearing eye-glasses), and also with classical observations that people can readily switch between different perturbation environments given enough experience (Martin et al., 1996a; Welch et al., 1993). It seems implausible that the ability to perform under such conditions could remain purely deliberate. A critical difference in these cases from those in which savings has been shown to be deliberate (Haith et al., 2015; Huberdeau et al., 2015b; Morehead et al., 2015) is that they involved repeated practice with the perturbed visual feedback. Theories of skill learning that posit that practice promotes a transition from deliberate processes to automaticity (Anderson, 1982; Ashby and Crossley, 2012; Fitts, Paul M., 1964) could therefore be the key to understanding savings. Specifically, might a deliberate re-aiming strategy become automatic with practice?

We sought to test the hypothesis that sustained practice would lead to savings being transformed from deliberate to automatic expression. Human participants adapted to a series of alternating perturbations over two days. It is known that deliberate re-aiming can be prohibited early in adaptation by limiting the amount of preparation time (PT) allowed prior to movement (Fernandez-Ruiz et al., 2011; Haith et al., 2015; Leow et al., 2017a, 2017b). We hypothesized that, if savings became supported by an automatic process, it would become expressible even when preparation time was limited.

An alternative way that practice might improve performance is if it leads to changes in the sensitivity to error of implicit recalibration (Gonzalez Castro et al., 2014; Herzfeld et al., 2014). We assessed potential changes in implicit recalibration using Aftereffect trials, which are uninfluenced by deliberate re-aiming (Benson et al., 2011; Morehead et al., 2015; Taylor et al., 2014; Werner et al., 2015). If a transformation occurs in the representation of savings from deliberate to automatic, then it should be evident even when preparation time is limited. In addition, the effect of limiting preparation time should dissociate from Aftereffect trials. If, instead, practice alters the error-sensitivity of implicit recalibration, then Aftereffect trials and Short-PT trials should remain congruent.

## Results

### Limited expression of learning in Short-PT and Aftereffect trials during the initial rotation cycle

Based on prior studies, we expected that limiting preparation time, as was imposed in Short-PT trials (Figure 1a), would lead to reduced expression of learning during the initial few trials of adaptation (Figure 1b-c) (Fernandez-Ruiz et al., 2011; Haith et al., 2015; Leow et al., 2017b). We also expected that explicitly instructing participants to withhold any deliberate re-aiming strategy (as was done for Aftereffect trials; Figure 1d) would have a similar effect (Benson et al., 2011; Morehead et al., 2015; Werner and Bock, 2010). This was indeed the case (Figure 2a & b). During the first few trials after onset of the rotation (trials 2-8), average reach angles were different across the three trial types (Long-PT trials, Short-PT trials, and Aftereffect trials; one-way ANOVA: F(2) = 3.55, p < 0.05). Post-hoc comparisons confirmed that compensation during this period was significantly less in Short-PT trials than in Long-PT trials (t = 4.31, p < 0.001), though the difference in reach direction between Aftereffect trials and Long-PT trials was not significant (t = 1.89, p = 0.077). Behavior did not differ significantly between the Short-PT and Aftereffect trials (t = 1.57, p = 0.14). This analysis therefore confirmed that limiting preparation time reduced the amount of compensation during early learning, but was inconclusive about whether behavior in Aftereffect trials differed from that in Long-PT trials.

**Figure 1:**
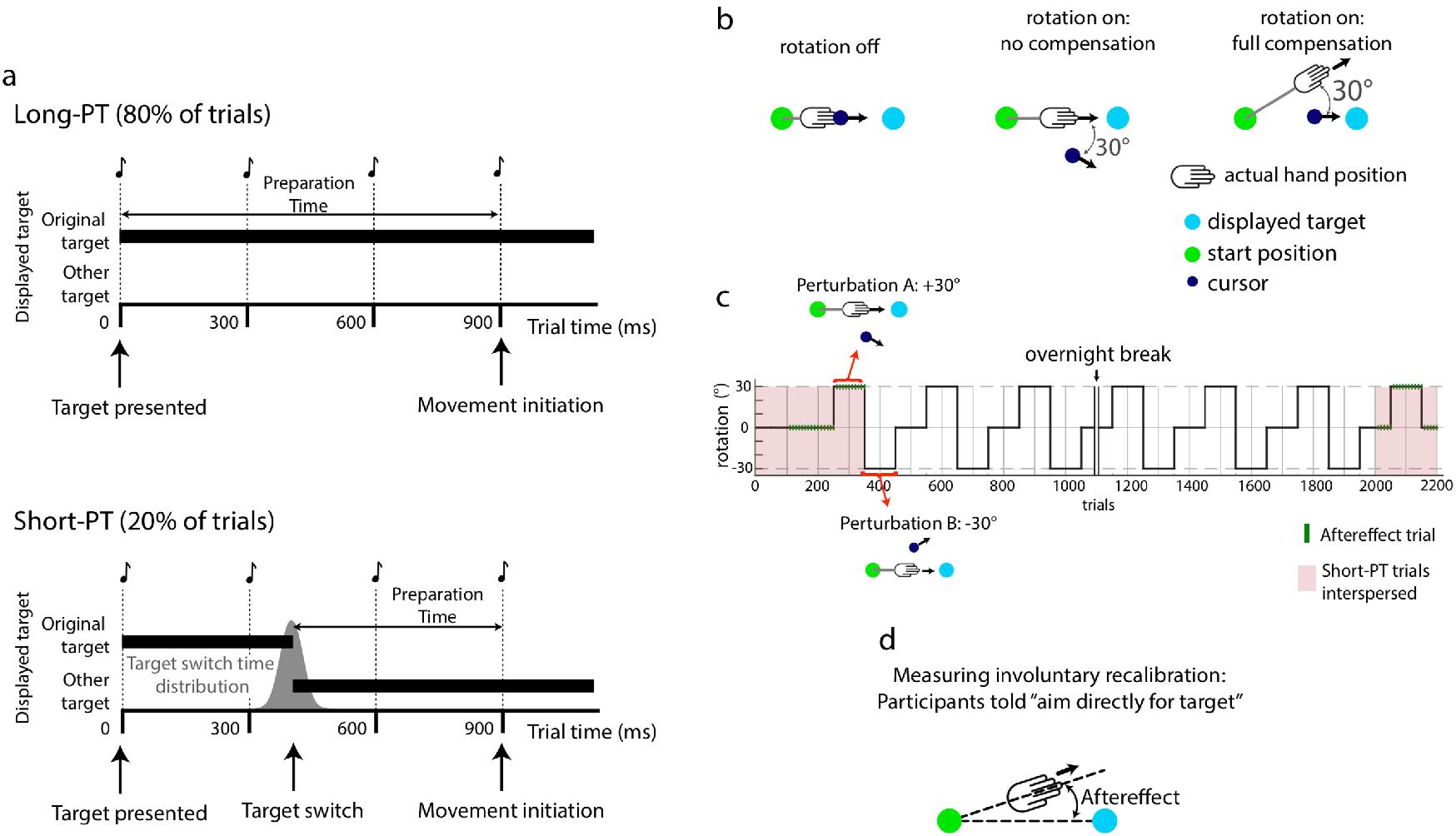
Experiment Design. Participants engaged in a reaching task. (a) Participants were required to initiate movement coincident with the fourth tone of a metronome. In the majority of trials (Long-PT trials, top panel), the target remained in place. In a subset of trials (Short-PT, lower panel), the target switched locations, from left to right or from right to left, just prior to the fourth beep, limiting allowed preparation time. (b) A rotation of the cursor path was imposed, requiring participants to adjust their movement direction relative to the target. (c) The direction of the rotation was varied in repeating cycles of two opposing rotation directions throughout a two-day experiment. (d) Aftereffect trials, in which participants were instructed to disengage any deliberate aiming strategy and “aim directly for the target”. Placement of Aftereffects trials within each cycle shown as green bars in (c).

**Figure 2:**
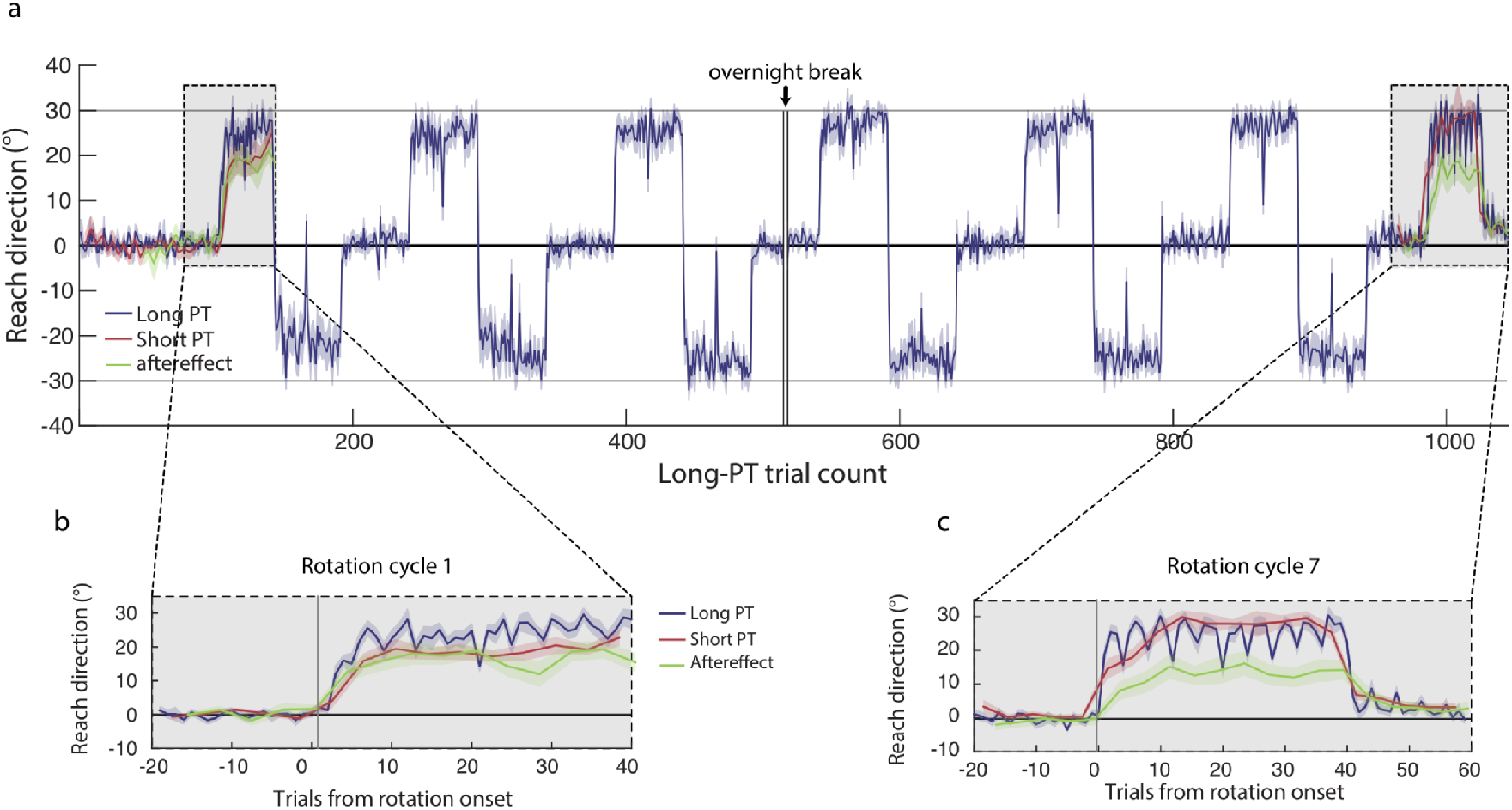
Experiment 1 Results. (a) Mean reach direction (+/- s.e.m.) across participants throughout the whole experiment. Blue: Long-PT trials; red: Short-PT trials; green: Aftereffect trials. Note that both Short-PT and Aftereffect trials were only present during the first and last cycle). (b) Enlarged view of mean behavior during Cycle 1. (c) Enlarged view of mean behavior during Cycle 7.

One problem with comparing learning across trial types within a fixed window of trials is that Aftereffect trials consistently occurred later in the block than the equivalent Short-PT trials. Thus, more learning was likely to have accrued prior to each Aftereffect trial than the other trial types, potentially masking a difference in behavior in these trials. We therefore conducted a finer-grained analysis that compared reach direction for each Short-PT or Aftereffect trial to the average reach direction for each of the two nearest-neighbor Long-PT trials (though excluding trials immediately following an Aftereffect trial; see below for justification as to why). This comparison reconfirmed the effect of limiting preparation time on extent of overall compensation (Short-PT vs Long-PT trials; t = 4.49, p < 0.001), and also revealed a significant difference in behavior between Aftereffect trials and regular, Long-PT trials (Aftereffect trials vs Long-PT trials; t = 5.42, p < 0.001).

We also examined behavior in each trial type at asymptote (trials 34 – 40). Here, there was also a significant difference among the trial types (one-way ANOVA: F(2) = 6.42, p < 0.01), with a significant difference between Long-PT and Aftereffect trials (t = 4.20, p < 0.001) and between Long-PT and Short-PT trials (t = 5.34, p < 0.001), but not between Aftereffect and Short-PT trials (t = 0.103, p = 0.92).

Thus, consistent with previous findings (Benson et al., 2011; Haith et al., 2015; Leow et al., 2017b; Morehead et al., 2015; Taylor et al., 2014), removing the influence of an aiming strategy during the first exposure to the perturbation, either by limiting preparation time (Short-PT trials) or by instruction (Aftereffect trials), significantly diminished the extent of compensation, particularly during early learning, confirming that participants compensated for the perturbation using a combination of deliberate re-aiming and implicit recalibration.

### Reversion toward baseline following Aftereffect trials

We noted that, in Long-PT trials that immediately followed Aftereffect trials, the reach direction was, on average, nearer to baseline compared to the Long-PT trial preceding the Aftereffect trial (Supplemental Figure 1; Cycle 1: t = 4.87, p < 0.001; Cycle 7: t = 5.21, p < 0.001). This effect was similar to those in previous studies showing that when adaptation is interrupted by a period of inactivity, the next reach following the interruption is closer to baseline than the reach prior to the interruption (Day and Singer, 1967; Hadjiosif and Smith, 2013; Morehead et al., 2015). Adaptation did, however, appear to recover rapidly in the subsequent trials. For this reason, post-Aftereffect trials (which, by design, were always Long-PT trials) were excluded from all analyses.

### Practice enabled savings for opposite rotations

During the course of the experiment, the direction of the cursor rotation periodically alternated between “rotation A” (30° or -30°, counterbalanced across participants), to “rotation B” (-30° or 30°), and finally to “null” (0°) (Figure 1c). Participants successfully adapted to both rotation directions (Figure 2a), and gradually improved the rate at which they adapted across cycles (rotation A, Cycle 7 vs. cycle 1: *t* = 4.24, *p* < 0.001; rotation B: *t* = 3.88, *p* < 0.01). Critically, this improvement was not attributable to differences in baseline at the start of each cycle, which was well-matched across cycles for all three trial types (First cycle vs. Last cycle; Long-PT: t = 0.687, p = 0.50; Short-PT: t = 0.0253, p = 0.98; and Aftereffect: t = 0.366, p = 0.72). Thus, following repeated exposures, participants were able to adapt to the alternating rotations and exhibited savings (faster learning) for both rotation directions when preparation time was long.

### The rate of implicit recalibration was not altered by practice

In order to determine which components of learning supported the savings seen following repeated practice, we re-introduced the probe trials (Aftereffect trials and Short-PT trials) in the final perturbation cycle (Figure 1c). Despite the strong savings seen in regular, Long-PT trials, performance in Aftereffect trials did not improve during the seventh cycle compared to the first cycle (Figure 3a & b; rate: t = 2.08, p = 0.052; asymptote: t = 2.48, p < 0.05). Implicit recalibration was actually marginally slower in the final cycle compared to the first. Thus, repeated practice had little effect on the rate of implicit recalibration and certainly could not account for the savings expressed in Long-PT trials.

**Figure 3:**
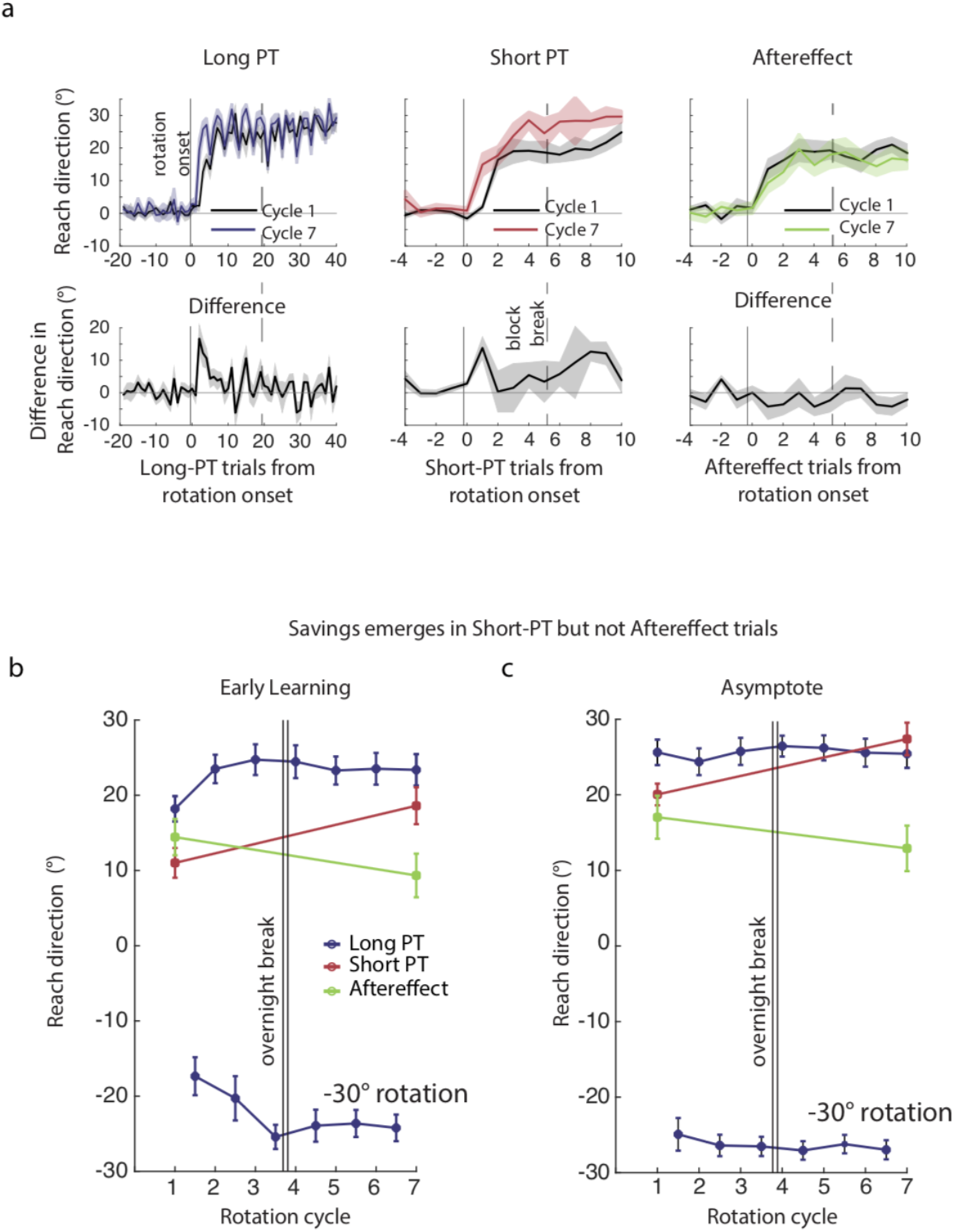
Effects of practice on rate of adaptation in Experiment 1. (a) Comparison of mean behavior across participants between Cycle 1 (gray) and Cycle 7 (colored) for each trial type. Lower set of panels in (a) show the mean difference in performance across cycles for each trial type. A positive difference indicates savings. (b) Extent of compensation early after perturbation onset (trials 2 – 8 following rotation onset) for each cycle. Lower blue line indicates early compensation during Long-PT trials following onset of perturbation B within each cycle. Behavior for Short-PT (red) and Aftereffect (green) trials was only measured in Cycles 1 and 7. (c) As (b), but showing asymptotic compensation (trials 32 – 40 following rotation onset) within each cycle.

By contrast, repeated practice significantly improved performance in trials in which preparation time was limited. The amount of learning expressed in Short-PT trials during initial adaptation to rotation A increased significantly from the first cycle to the seventh cycle (rate: t = 2.84, p < 0.05; asymptote: t = 5.23, p < 0.001). There was also a significant interaction among the three trial types for adaptation rate (Figure 3b; 2-way ANOVA interaction: F(2) = 1.85, p < 0.05).

In summary, we found that the implicit recalibration did not change despite extensive practice adapting to a perturbation. Practice did, however, lead to a qualitative change in the nature of the memory for adaptation, apparent in the fact that participants were able to express more of their learning when preparation time was limited, suggesting a transition from a computationally expensive deliberate process to one that was automatic.

### Savings under limited preparation time emerged gradually with practice

The results of Experiment 1 suggested that repeated exposure to a perturbation led to a qualitative change in behavior, with faster, more automatic compensation. However, because we only probed during the first and the final cycles, we were unable to determine the time-course over which this change occurred. We therefore conducted another experiment, Experiment 2, in which Short-PT trials were included throughout the experiment in order to determine the time-course over which this transition occurred (Figure 4a).

Behavior during the first rotation cycle paralleled that of Experiment 1. Limiting preparation time led to a significant reduction in compensation during the first few trials following rotation onset (Figure 4b; t = 4.17, p < 0.001). When the imposed perturbation switched direction (from perturbation A to B), compensation in Short-PT trials continued to lag that in Long-PT trials (t = 3.84, p < 0.01).

**Figure 4:**
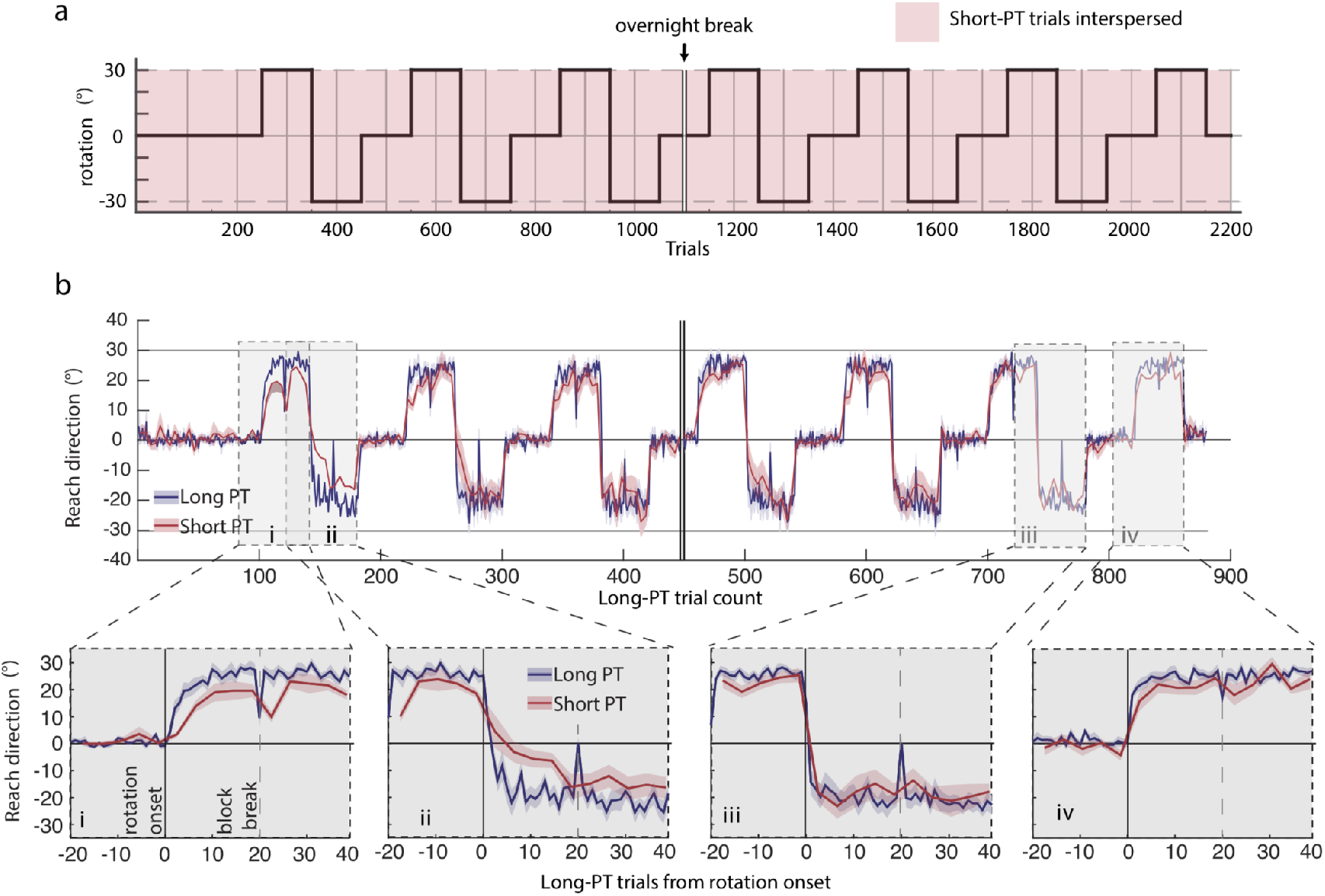
Results for Experiment 2. (a) Participants adapted to the same repeating set of rotations as in Experiment 1, but with Long- and Short-PT trials included throughout all cycles (and no Aftereffect trials) (b). Average behavior in Long-PT (blue) and Short-PT (red) trials through the whole experiment. Lower panels show enlarged view of behavior during (i) the onset of perturbation A during Cycle 1, (ii) the transition from perturbation A to perturbation B during Cycle 1 (iii), the last transition from perturbation A and perturbation B, which occurred in Cycle 6 (iv), and the onset of perturbation A in Cycle 7.

As in Experiment 1, Long-PT trials exhibited savings between the first and seventh cycles of rotation A (t = 4.95, p < 0.001). A similar savings effect was evident in Short-PT trials (Figure 5a; t = 3.10, p < 0.05). To determine whether the development of savings expressed in Long-PT and Short-PT trials followed a similar time-course across the seven cycles of the rotation, we conducted a linear two-way mixed-effects analysis. This analysis showed no interaction affect between cycle and trial type, suggesting that the development of savings followed a similar time-course in the Long-PT and Short-PT trials (Figure 5b & c; cycle-by-trial type interaction for rate: X^2^(1) = 1.95, p = 0.16; and for asymptote: X^2^(1) = 0.27, p = 0.60).

**Figure 5:**
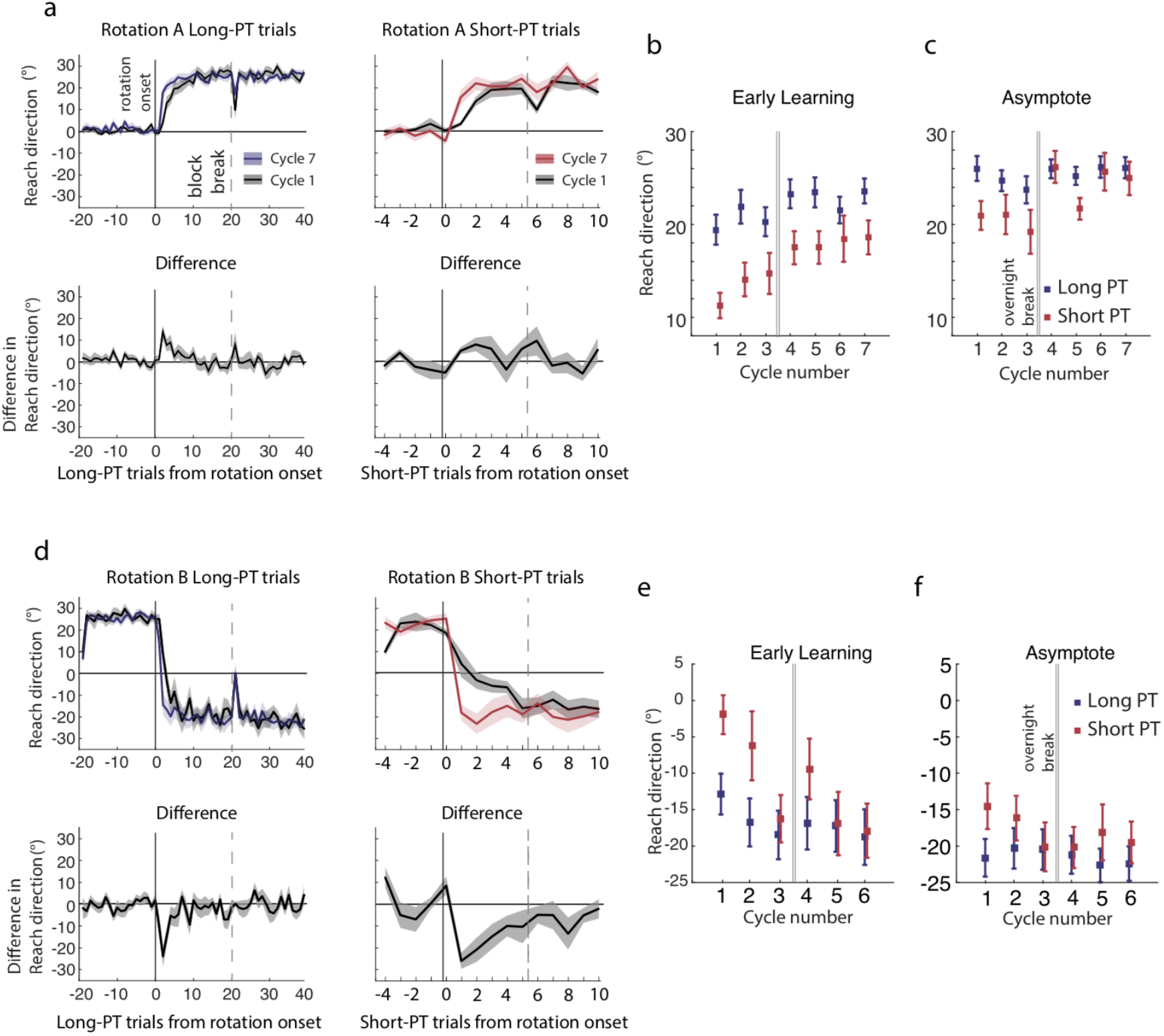
Effects of practice on the time course of adaptation in Experiment 2. (a) Mean behavior across participants within Cycle 1 (gray) and Cycle 7 (color) for both trial types. Lower panels show mean difference in performance across cycles. (b) Mean performance early after onset of perturbation A (trials 2-8), across cycles for Long-PT (blue) and Short-PT (red) trials. (c), as (b) but for mean performance at asymptote (trials 32-40) under perturbation A within each cycle. (d-f), as (a-c), but showing behavior under perturbation B within Cycles 1 and 6 (d) and across cycles (e,f).

Short-PT probe trials following the transition to rotation B revealed an even more dramatic savings effect from practice across cycles of the rotation (Figure 5d). In Long-PT trials, participants exhibited modest improvements in their rate of adaptation to perturbation B (Figure 5e; t = 3.21, p < 0.01), exactly as we had found in Experiment 1. In Short-PT trials, by contrast, participants struggled to compensate during the first perturbation cycle, but by the final cycle the response to the rotation was not detectably different from Long-PT trials (Figure 5e; t = 0.82, p = 0.42). A mixed effects model found a significant interaction between cycle and trial type for rotation B over the first few trials (Figure 5e; chi-s(1) = 13.7, p < 0.001), although not significantly for asymptote (Figure 5f; chi-s(1) = 1.85, p = 0.17).

In summary, the results of Experiment 2 show that the improvements in performance during Short-PT trials emerged gradually with practice and, at least for rotation B, differently from Long-PT trials.

## Methods

### Experiment Participants

41 right-handed participants with no known neurological impairments took part in this study (18 – 40 years old, 25 women). The study was approved by the Johns Hopkins University School of Medicine Institutional Review Board.

### Experimental Setup

Participants were seated at a glass-surfaced table with their right forearm supported by a splint that allowed nearly frictionless planar arm movement. Participants’ arms were obstructed from their own view by a mirror, on which was projected a display from a downward-facing LCD monitor installed above the mirror (60 Hz refresh rate; LG).

Each participant’s hand position was recorded by a Flock of Birds magnetic sensor (130 Hz; Ascension Inc., Shelburne, VT) placed under each participant’s index finger. Hand position was reported to participants in near real-time via a cursor (a filled blue circle, diameter 0.5 cm) displayed on the screen. Visual feedback of the cursor had a delay of approximately 100 ms (40 ms delay in the Flock of Birds and an approximately 60 ms delay in the visual display).

### Experiment 1

21 participants took part in Experiment 1, although two were excluded from analysis because more than 50% of their Short-PT trials were directed towards the wrong target. Participants made rapid “shooting” movements using their right arm from a central start location (a solid green circle, diameter 1 cm) through a target (a solid light-blue circle, diameter 1 cm). The target could appear at one of two locations, positioned 8 cm either to the right or left of the start location. Participants were trained to initiate their reaching movement coincident with the fourth of four audible tones (Figure 1a). The tone sequence began 200 ms following steady placement of the cursor inside the start marker. Successive tones were played at intervals of 300 ms. On each trial, one of the two targets was presented at the onset of the first tone. During Long-PT trials, the initially-presented target remained on the screen either until the participant reached 9 cm radially from the start position, or 2.5 s passed from the time of the first tone. For Short-PT trials, the target abruptly switched sides at a variable time prior to the fourth ring tone (Figure 1a), and remained there for the duration of the trial as with Long-PT trials.

A visuomotor perturbation in the form of a ±30° rotation of the path of the cursor about the start position (Figure 1b) was applied to movements directed towards the right half of the workspace in repeating cycles throughout the experiment (Figure 1c). Leftward-directed movements had no rotation at any time throughout the experiment. Trials with rightward-directed movements and trials with leftward-directed movements were pseudorandomly interleaved throughout each cycle. Each cycle included 150 trials of rightward-directed, and 150 trials of leftward-directed, movements. The rotation schedule was 50 trials of null rotation, 50 trials of rotation A, and 50 trials of rotation B. The direction of the cursor rotation under rotation A and B was counter-balanced across participants in the experiment. 11 participants had rotation A as a clockwise rotation, and 10 participants had it as a counterclockwise rotation. Seven cycles were included across the duration of the experiment. The seventh cycle omitted rotation B. The experiment was divided into blocks of 100 total trials (grey vertical lines in Figure 1c), with brief breaks in between blocks, plus an overnight break between cycles 3 and 4. Changes in the rotation occurred in the middle of blocks.

We included three different types of trials to probe participants’ mode of compensation for the perturbation. The majority of trials were designated as long-preparation-time (Long-PT) trials. In these trials, the target remained in its original location for the duration of the trial so that participants had 1.2 seconds to prepare their movement. During short-preparation time (Short-PT) trials, the target location abruptly switched to the opposite possible target position prior to the fourth tone (Figure 1a). The time at which the target switched locations was randomized for each Short-PT trial by sampling from a Gaussian distribution with a mean of 300 ms and a standard deviation of 25 ms. Short-PT trials were included among the more frequent Long-PT trials only during the first rotation and during the seventh and final rotation (Figure 1c). Within blocks where they were present, Short-PT trials were randomly interspersed among Long-PT trials such that for every 10 total trials, two were Short-PT (one to each target) and eight were Long-PT (four to each target). No Short-PT trials were permitted as the first or last trial in each sequence of 10 trials.

We also included an additional, Aftereffect probe (Figure 1d), designed to directly assess implicit recalibration. Prior to these probes, participants were explicitly instructed to aim directly for the presented target, rather than applying a strategy or deliberately aiming in a direction other than towards the target (Benson et al., 2011; Day et al., 2016; Morehead et al., 2015; Werner et al., 2015). Prior to these trials, text appeared on the participants’ screen for 4.5 seconds reading: “On the next trial / take your time / and aim directly for the target”. All participants were literate in English. Participants were also verbally instructed at the beginning of each session of the experiment that during these Aftereffect probe trials, no cursor would be visible, no audible tone sequence would sound, no movement initiation time constraints were in place, and they were to reach for the target as if they wanted their finger to intersect with the target.

A pair of Aftereffect probes, one for each target direction, followed each series of 10 Long- or Short-PT trials in blocks when they were present (Figure 1c). In Experiment 1, Aftereffect probes were included in all blocks for which Short-PT trials were present, except for the initial familiarization block (Figure 1c).

Participants were instructed that for Long- and Short-PT trial types they were to prioritize the timing of their movement initiation. They were instructed to be as accurate as possible in hitting the target with the cursor, and to reach with a consistent, fast speed (between 4.5 cm/s and 13 cm/s). Feedback regarding movement timing and movement speed was provided following every Long- and Short-PT trial through visual displays on the screen (similar to (Haith et al., 2015)).

Cursor feedback during the movement was provided throughout each Long- and Short-PT trial. The cursor disappeared once participants reached 9 cm radially from the start position. The cursor was not visible during the return movement, until the participants’ hand was within 2 cm of the start position. Any cursor manipulations (i.e. the rotations) were turned off during the inter-trial period. During Aftereffect probes, no cursor feedback was provided apart from during return movements.

### Experiment 2

20 participants took part in Experiment 2, and 3 were excluded because 50% of their Short-PT trials were directed towards the wrong target. The reaching task and rotation schedule remained the same for Experiment 2 as in Experiment 1. Experiment 2 included Short-PT trials throughout the entire experiment, rather than just the first and final rotation cycles as had been the case for Experiment 1. No Aftereffect trials were included in Experiment 2. Experiment 2, like Experiment 1, was conducted in two sessions across two consecutive days.

### Data analysis

All data were analyzed offline in Matlab (The Mathworks, Natick, MA) and in R (The R Project, www.r-project.org). Kinematic data were smoothed with a 2^nd^-order Savitzky-Golay interpolation filter with half width 54 ms. These smoothed signals were then differentiated to obtain velocity. The time of movement initiation was determined by searching from the peak velocity backwards in time to find the last time at which tangential velocity exceeded a threshold of 2 cm/s. Reach direction was determined by computing the angle of the instantaneous velocity at 100 ms after movement onset. Trials during which participants either failed to reach or abruptly altered their initial reach direction after having reached 2 cm from the start position were excluded from analysis (on average, 5 trials were excluded per participant for this reason). This type of error was most likely to have occurred during Short-PT trials, due to participants initially moving towards the original target location. Participants were excluded from further analysis if fewer than 50% of their Short-PT trials were directed towards the correct target.

The initial learning rate during a given rotation cycle was quantified as the average compensation over the first few trials of that cycle. Following a similar approach as in (Haith et al., 2015), we assessed initial learning during Long-PT trials based on the mean reach direction over the initial eight Long-PT trials, though we exclude the first trial following rotation onset and any post-Aftereffect trials from this average. For Short-PT trials and Aftereffect probes, the average reach direction in the initial two trials of each type following rotation onset was taken as the initial learning measure. Similarly, the final eight trials (for Long-PT trials), and final two trials (for Short-PT and Aftereffect trials), in each rotation were averaged and used as a summary measure for asymptotic behavior (excluding post-Aftereffect trials).

For Experiment 1, a 2-way analysis of variance (ANOVA) test was conducted on the early adaptation measure, with trial type (Long PT, Short PT, and Aftereffect) and rotation cycle used as factors. In the event of an interaction between trial type and cycle, t-tests were planned to detect any difference among groups in early learning or asymptote during the first and the final rotation cycles, and to test for savings from the first to the final rotation cycle for each trial type. A linear mixed-effects model analysis was conducted for Experiment 2, using trial type (Long- and Short-PT), and cycle (cycles 1 to 6) as fixed effects, and subject as a random factor.

## Discussion

Our experiments showed that repeated exposure to a pair of alternating rotations led to a qualitative change in the ability to express memory for adaptation. We measured the extent of implicit recalibration using Aftereffect trials and found, consistent with previous work, that implicit recalibration accounted for only a fraction of overall learning, implying the existence of additional re-aiming processes (Morehead et al., 2015; Taylor et al., 2014). Importantly, the rate of implicit recalibration remained invariant despite multiple exposures to the perturbation and clear savings in regular, Long-PT trials. This finding refutes suggestions that savings might be attributable to modulation of the sensitivity of implicit recalibration (Herzfeld et al., 2014).

The invariance of implicit recalibration despite practice was in contrast to what we observed in Short-PT trials in which preparation time was limited. During the first cycle, limiting preparation time reduced overall compensation to an extent comparable to Aftereffect trials. This is consistent with the idea that limiting preparation time prohibited the use of a deliberate re-aiming strategy (Fernandez-Ruiz et al., 2011; Haith et al., 2015; Leow et al., 2017b). However, the effect of limiting preparation time diminished with practice. Participants became able to express the bulk of their learning regardless of allowed preparation time. Since this practice effect was not due to improved implicit recalibration, we conclude that it was attributable to participants becoming able to express their learning more rapidly and automatically.

This transformation is consistent with more general theories of motor skill learning that posit a transition from deliberate to automatic modes of control following practice (Anderson, 1982; Ashby and Crossley, 2012; Economides et al., 2015; Fitts and Posner; Fitts, Paul M., 1964; Honda et al., 1998; Moors and De Houwer, 2006). The transition from deliberate to automatic control is more usually established through dual-task paradigms, which allow the reliance on cognitive resources to be measured at different points during learning (Schneider and Shiffrin, 1977). In addition to a reduction in cognitive load, automaticity has also been characterized in terms of processing speed, and whether or not responses are obligatory (Cohen et al., 1992; Logan, 1980). These other facets of automatic behavior have been relatively little studied in comparison to effects associated with cognitive load, and it is unclear exactly how these effects of practice are inter-related. We suggest that limiting response times might offer an alternative, perhaps more powerful approach to investigate these effects. Restricting preparation time has been demonstrated to limit deliberative reasoning in more abstract decision-making tasks (Keramati et al., 2011). We have also recently used a similar approach to establish that learned motor responses become faster and become habitual through practice (Hardwick et al., 2017). Future work will be necessary to fully understand the relationship between processing time, cognitive load, and whether or not responses are habitual.

### Overcoming interference through automaticity

A key difference from our previous work exploring the influence of movement preparation time on the expression of savings (Haith et al., 2015) is in the way we washed out participants’ learning in between exposures to the perturbation(s). Previously, we washed out participants by switching off the perturbation for 20 trials and allowing participants an overnight break. In this set of experiments, we washed participants out by imposing a counter-perturbation followed by no perturbation. We found this be an effective means of ensuring that behavior returned to a fixed baseline in all trial types, allowing us to directly compare the response to the introduction of the perturbation across cycles.

Imposing a counter-perturbation (perturbation B) also created the possibility of interference between memories for the two perturbations (Krakauer et al., 2005). Indeed, we found that the extent of savings in regular, Long-PT trials was weaker than is typical in adaptation experiments, suggesting partial interference. This interference was fully overcome with practice alternating between different perturbations, as would be expected based on classical experiments on adapting to alternating visual shifts induced by prisms (Martin et al., 1996b). Experiment 2 revealed that the ability to overcome interference became established over roughly the same time course as the emergence of savings in Short-PT trials, suggesting that automatization might have been critical to the ability to overcome interference effects. This is consistent with the suggestion that interference might be a cognitive phenomenon, brought about by blocking retrieval of a learned compensation strategy (Krakauer et al., 2005; Yin and Wei, 2014). Automatized compensation strategies might be less vulnerable to such cognitive interference effects than deliberate strategies, hence enabling savings for both rotation directions.

### If skill learning is initially declarative, how can patients with amnesia learn new motor skills?

Perhaps no experimental result has influenced motor learning theory more than that of patient H.M: despite severe anterograde and retrograde amnesia, H.M. and other patients like him were capable of learning novel motor abilities like mirror drawing, despite having no recollection of ever having done the practiced tasks (Cohen and Squire, 1980; Milner, 1962). These findings directly gave rise to the deeply embedded notion that motor skills are procedural, and distinct from declarative memory systems (Cohen and Squire, 1980). How can the H.M. result be reconciled with our model of a transformation from deliberate to automatic control? The answer, we suggest, is that the processes needed for deliberate control are in fact intact in amnesic patients (Schacter et al., 1982; Squire and Zola, 1998; Tulving, 1985), even though the ability to build long-term memories for these processes is impaired. Amnesic patients are unimpaired at most cognitive tasks and basic reasoning abilities (Schacter et al., 1982, 1982; Squire and Zola, 1998; Tulving, 1985), provided the tasks do not require holding specific facts in memory beyond the capacity of their short-term memory. H.M. could have been able to gradually learn new skills by rapidly automatizing fragments of the skill within each session. These automatized fragments *could* then have been retrieved in subsequent sessions, leaving less work for deliberate control. Iterating this fragmentary automatization and retrieval would ultimately allow a new, deliberate skill to be gradually acquired and retained across sessions, even though the skill initially depended on declarative processes.

### Relationship between adaptation and motor skill learning

Our findings help to clarify the relationship between adaptation and motor skill learning. Adaptation paradigms, along with other similar cerebellum-dependent forms of motor learning, like smooth pursuit (Yang and Lisberger, 2014), are often considered to represent models of motor learning in a general sense. However, the relationship of simple adaptation tasks to more complex real-world skills is questionable (Krakauer and Mazzoni, 2011; Wulf and Shea, 2002). Adaptation occurs in minutes, whereas real-world skills are learned over days, weeks or months. Adaptation tasks can be solved perfectly through instruction (Mazzoni and Krakauer, 2006), unlike real-world skills where extensive practice is typically necessary even if the required actions are easily communicable. Furthermore, adaptation is a transient state that, unlike other skills, tends to deteriorate rather than consolidate with the passage of time (Kitago et al., 2013).

The relevance of adaptation to motor skill learning hinges on the phenomenon of savings, which represents the only real long-term memory associated with adaptation. Recent results showing that savings is largely attributable to retrieval of deliberate aiming strategies (Haith et al., 2015; Morehead et al., 2015) therefore cause significant difficulty for the idea that adaptation might in some way model the acquisition of more general motor skills. Our findings offer a potential reconciliation, showing that a compensatory strategy that is initially applied deliberately can be applied automatically following practice. This transition mirrors the transition from declarative to procedural memory that has commonly been invoked in theories of skill learning (Anderson, 1982; Fitts and Posner). Thus, in a restricted sense, adaptation paradigms do encompass a model of more general skill learning processes. Nevertheless, the presence of implicit recalibration, which appears to be insensitive to practice-related effects, actually significantly complicates behavior in such paradigms. We suggest that skill learning might be better studied in paradigms that more effectively isolate the deliberate-to-automatic transition.

## Acknowledgements

Thanks to members of the BLAM lab for helpful criticism and discussion. We thank Aaron Wong, Alex Forrence, Reza Shadmehr, and Amy Bastian.

---

**Supplemental Figure 1:**
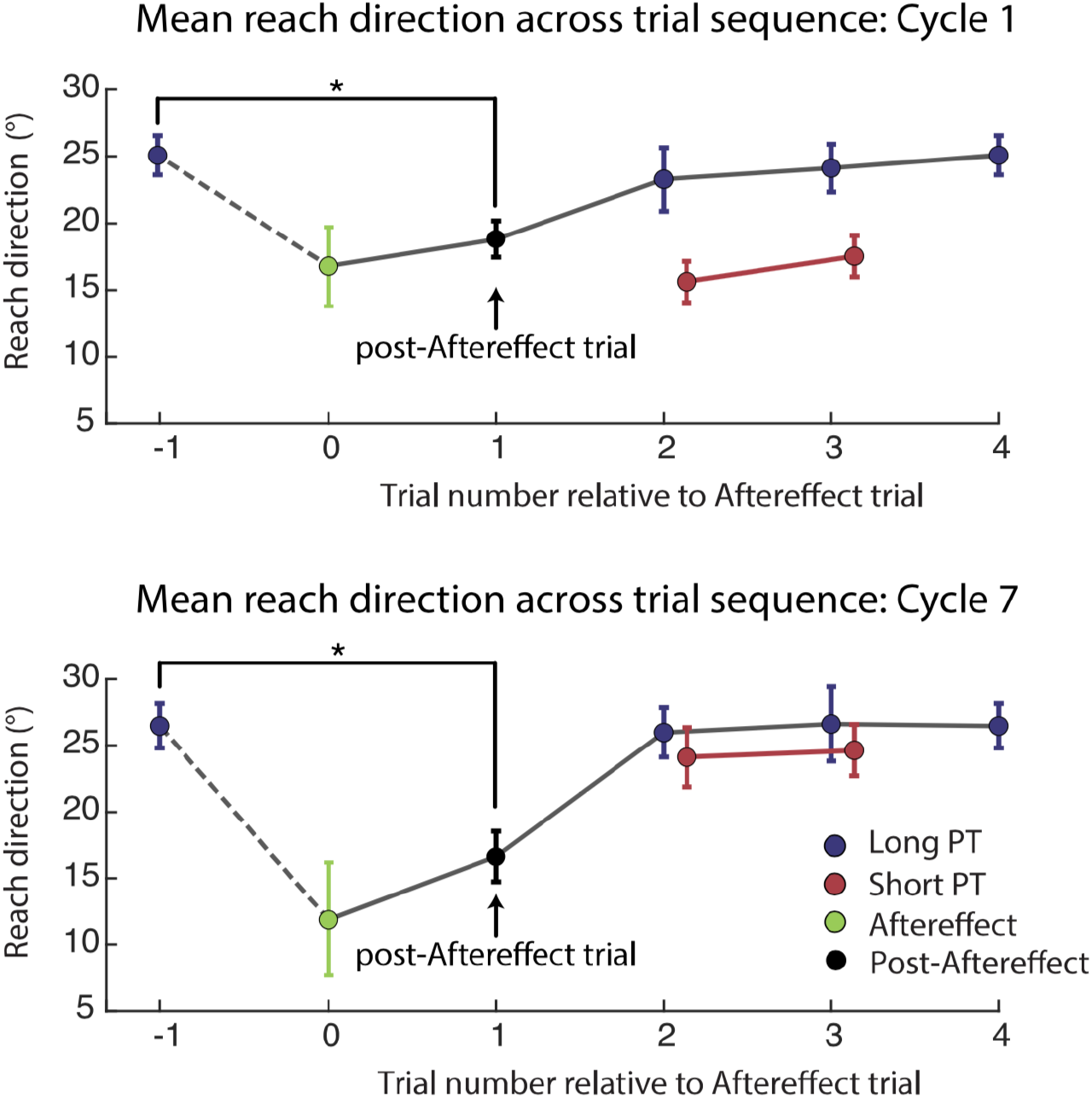
Behavior following aftereffect trials. Top panel: Mean performance surrounding Aftereffect trial (trial 0). Black cirles represent Long-PT trials that occurred immediately after the Aftereffect trial. Blue circles represent mean performance in Long-PT trials at other positions relative to the Aftereffect trial. Red circles represent mean performance in the first Short-PT trial after each Aftereffect trial (which occurred either 2 or 3 trials later). Bottom Panel: As Top Panel, but for Cycle 7. *indicates p < 0.05

## References

Anderson, J. (1982). Acquisition of cognitive skill. Psychol. Rev. 89.

Ashby, F.G., and Crossley, M.J. (2012). Automaticity and multiple memory systems. Wiley Interdiscip. Rev. Cogn. Sci. 3, 363–376.

Benson, B.L., Anguera, J.A., and Seidler, R.D. (2011). A spatial explicit strategy reduces error but interferes with sensorimotor adaptation. J. Neurophysiol. 105, 2843–2851.

Brashers-Krug, T., Shadmehr, R., and Bizzi, E. (1996). Consolidation in human motor memory. Nature 382, 252–255.

Cohen, N.J., and Squire, L.R. (1980). Preserved learning and retention of pattern-analyzing skill in amnesia: dissociation of knowing how and knowing that. Science 210, 207–210.

Cohen, J.D., Servan-Schreiber, D., and McClelland, J.L. (1992). A Parallel Distributed Processing Approach to Automaticity. Am. J. Psychol. 105, 239–269.

Cunningham, H.A. (1989). Aiming error under transformed spatial mappings suggests a structure for visual-motor maps. J. Exp. Psychol. Hum. Percept. Perform. 15, 493.

Day, H.R., and Singer, G. (1967). Sensory adaptation and behavioral compensation with spatially transformed vision and hearing. Psychol. Bull. 67, 307–322.

Day, K.A., Roemmich, R.T., Taylor, J.A., and Bastian, A.J. (2016). Visuomotor Learning Generalizes Around the Intended Movement. eNeuro 3, ENEURO.0005-16.2016.

Ebbinghaus, H. (1913). Memory (HA Ruger & CE Bussenius, Trans.). N. Y. Columbia Univ. Teach. Coll. Work Publ. 1885.

Economides, M., Kurth-Nelson, Z., Lübbert, A., Guitart-Masip, M., and Dolan, R.J. (2015). Model-Based Reasoning in Humans Becomes Automatic with Training. PLOS Comput. Biol. 11, e1004463.

Fernandez-Ruiz, J., Wong, W., Armstrong, I.T., and Flanagan, J.R. (2011). Relation between reaction time and reach errors during visuomotor adaptation. Behav. Brain Res. 219.

Fitts, P., and Posner, M. Human performance.

Fitts, Paul M. (1964). Perceptual Motor Skill Learning. In Categories of Human Learning, A.W. Melton, ed. (New York, NY: Academic Press Inc.), pp. 244–283.

Gonzalez Castro, L.N., Hadjiosif, A.M., Hemphill, M.A., and Smith, M.A. (2014). Environmental consistency determines the rate of motor adaptation. Curr. Biol. CB 24, 1050–1061.

Hadjiosif, A., and Smith, M.A. (2013). Savings is restricted to the temporally labile component of motor adaptation.

Haith, A.M., and Krakauer, J.W. (2014). Motor Learning: The Great Rate Debate. Curr. Biol. 24, R386–R388.

Haith, A.M., Huberdeau, D.M., and Krakauer, J.W. (2015). The Influence of Movement Preparation Time on the Expression of Visuomotor Learning and Savings. J. Neurosci. 35, 5109–5117.

Hardwick, R.M., Forrence, A.D., Krakauer, J.W., and Haith, A.M. (2017). Skill Acquisition and Habit Formation as Distinct Effects of Practice. bioRxiv 201095.

Herzfeld, D.J., Vaswani, P.A., Marko, M.K., and Shadmehr, R. (2014). A memory of errors in sensorimotor learning. Science 345, 1349–1353.

Honda, M., Deiber, M.P., Ibáñez, V., Pascual-Leone, A., Zhuang, P., and Hallett, M. (1998). Dynamic cortical involvement in implicit and explicit motor sequence learning. A PET study. Brain 121, 2159–2173.

Huang, V.S., Haith, A., Mazzoni, P., and Krakauer, J.W. (2011). Rethinking motor learning and savings in adaptation paradigms: model-free memory for successful actions combines with internal models. Neuron 70.

Huberdeau, D.M., Krakauer, J.W., and Haith, A.M. (2015a). Dual-process decomposition in human sensorimotor adaptation. Curr. Opin. Neurobiol. 33, 71–77.

Huberdeau, D.M., Haith, A.M., and Krakauer, J.W. (2015b). Formation of a long-term memory for visuomotor adaptation following only a few trials of practice. J. Neurophysiol. 114, 969–977.

Joiner, W.M., and Smith, M.A. (2008). Long-term retention explained by a model of short-term learning in the adaptive control of reaching. J. Neurophysiol. 100, 2948–2955.

Keramati, M., Dezfouli, A., and Piray, P. (2011). Speed/Accuracy Trade-Off between the Habitual and the Goal-Directed Processes. PLOS Comput. Biol. 7, e1002055.

Kitago, T., Ryan, S.L., Mazzoni, P., Krakauer, J.W., and Haith, A.M. (2013). Unlearning versus savings in visuomotor adaptation: comparing effects of washout, passage of time, and removal of errors on motor memory. Front. Hum. Neurosci. 7, 307.

Kluzik, J., Diedrichsen, J., Shadmehr, R., and Bastian, A.J. (2008). Reach Adaptation: What Determines Whether We Learn an Internal Model of the Tool or Adapt the Model of Our Arm? J. Neurophysiol. 100, 1455–1464.

Kojima, Y., Iwamoto, Y., and Yoshida, K. (2004). Memory of learning facilitates saccadic adaptation in the monkey. J. Neurosci. Off. J. Soc. Neurosci. 24, 7531–7539.

Krakauer, J.W., and Mazzoni, P. (2011). Human sensorimotor learning: adaptation, skill, and beyond. Curr. Opin. Neurobiol. 21, 636–644.

Krakauer, J.W., Ghilardi, M.-F., and Ghez, C. (1999). Independent learning of internal models for kinematic and dynamic control of reaching. Nat. Neurosci. 2, 1026–1031.

Krakauer, J.W., Pine, Z.M., Ghilardi, M.-F., and Ghez, C. (2000). Learning of visuomotor transformations for vectorial planning of reaching trajectories. J. Neurosci. 20, 8916–8924.

Krakauer, J.W., Ghez, C., and Ghilardi, M.F. (2005). Adaptation to visuomotor transformations: consolidation, interference, and forgetting. J. Neurosci. Off. J. Soc. Neurosci. 25, 473–478.

Lackner, J.R., and Lobovits, D. (1977). Adaptation to displaced vision: evidence for prolonged after-effects. Q. J. Exp. Psychol. 29, 65–69.

Leow, L.-A., Marinovic, W., Riek, S., and Carroll, T.J. (2017a). Cerebellar anodal tDCS increases implicit learning when strategic re-aiming is suppressed in sensorimotor adaptation. PLOS ONE 12, e0179977.

Leow, L.-A., Gunn, R., Marinovic, W., and Carroll, T.J. (2017b). Estimating the implicit component of visuomotor rotation learning by constraining movement preparation time. J. Neurophysiol. jn.00834.2016.

Logan, G.D. (1980). Attention and automaticity in Stroop and priming tasks: Theory and data. Cognit. Psychol. 12, 523–553.

Martin, T.A., Keating, J.G., Goodkin, H.P., Bastian, A.J., and Thach, W.T. (1996a). Throwing while looking through prisms. Brain 119, 1199–1211.

Martin, T.A., Keating, J.G., Goodkin, H.P., Bastian, A.J., and Thach, W.T. (1996b). Throwing while looking through prisms. I. Focal olivocerebellar lesions impair adaptation. Brain J. Neurol. 119 (Pt 4), 1183–1198.

Mazzoni, P., and Krakauer, J.W. (2006). An implicit plan overrides an explicit strategy during visuomotor adaptation. J. Neurosci. 26, 3642–3645.

Milner, B. (1962). Les troubles de la memoire accompagnant des lesions hippocampiques bilaterales. In Physiologie de L’hippocampe, (Paris: Centre National de la Recherche Scientifique), pp. 257–272.

Moors, A., and De Houwer, J. (2006). Automaticity: a theoretical and conceptual analysis. Psychol. Bull. 132, 297–326.

Morehead, J.R., Qasim, S.E., Crossley, M.J., and Ivry, R. (2015). Savings upon Re-Aiming in Visuomotor Adaptation. J. Neurosci. 35, 14386–14396.

Schacter, D.L., Wang, P.L., Tulving, E., and Freedman, M. (1982). Functional retrograde amnesia: A quantitative case study. Neuropsychologia 20, 523–532.

Schneider, W., and Shiffrin, R. (1977). Controlled and automatic human information processing: I. Detection, search, and attention. Psychol. Rev. 84, 1–66.

Shadmehr, R., and Mussa-Ivaldi, F.A. (1994). Adaptive representation of dynamics during learning of a motor task. J. Neurosci. 14, 3208–3224.

Squire, L.R., and Zola, S.M. (1998). Episodic memory, semantic memory, and amnesia. Hippocampus 8, 205–211.

Taylor, J.A., Klemfuss, N.M., and Ivry, R.B. (2010). An Explicit Strategy Prevails When the Cerebellum Fails to Compute Movement Errors. Cerebellum Lond. Engl. 9, 580–586.

Taylor, J.A., Krakauer, J.W., and Ivry, R.B. (2014). Explicit and Implicit Contributions to Learning in a Sensorimotor Adaptation Task. J. Neurosci. 34, 3023–3032.

Tulving, E. (1985). Memory and consciousness. Can. Psychol. Can. 26, 1–12.

Villalta, J.I., Landi, S.M., Flo, A., and Della-Maggiore, V. (2013). Extinction Interferes with the Retrieval of Visuomotor Memories Through a Mechanism Involving the Sensorimotor Cortex. Cereb. Cortex N. Y. N 1991.

Welch, R.B., Bridgeman, B., Anand, S., and Browman, K.E. (1993). Alternating prism exposure causes dual adaptation and generalization to a novel displacement. Percept. Psychophys. 54, 195–204.

Werner, S., and Bock, O. (2010). Mechanisms for visuomotor adaptation to left–right reversed vision. Hum. Mov. Sci. 29, 172–178.

Werner, S., Aken, B.C. van, Hulst, T., Frens, M.A., Geest, J.N. van der, Strüder, H.K., and Donchin, O. (2015). Awareness of Sensorimotor Adaptation to Visual Rotations of Different Size. PLOS ONE 10, e0123321.

Wolpert, D.M., Ghahramani, Z., and Jordan, M.I. (1995). An internal model for sensorimotor integration. Science 269, 1880–1882.

Wulf, G., and Shea, C.H. (2002). Principles derived from the study of simple skills do not generalize to complex skill learning. Psychon. Bull. Rev. 9, 185–211.

Yang, Y., and Lisberger, S.G. (2014). Role of plasticity at different sites across the time course of cerebellar motor learning. J. Neurosci. Off. J. Soc. Neurosci. 34, 7077–7090.

Yin, C., and Wei, K. (2014). Interference from mere thinking: mental rehearsal temporarily disrupts recall of motor memory. J. Neurophysiol. 112, 594–602.

Zarahn, E., Weston, G.D., Liang, J., Mazzoni, P., and Krakauer, J.W. (2008). Explaining savings for visuomotor adaptation: linear time-invariant state-space models are not sufficient. J. Neurophysiol. 100, 2537–2548.

